# DNA methylation oscillation defines classes of enhancers

**DOI:** 10.1101/262212

**Authors:** Emanuele Libertini, Rifat A. Hamoudi, Simon Heath, Lee Lancashire, Arcadio Rubio Garcia, Luigi Grassi, Kate Downes, Willem H. Ouwehand, Biola-Maria Javierre, Jonathan Cairns, Steven Wingett, Dirk S. Paul, Marta Gut, Ivo G. Gut, Joost H. A. Martens, Alexandr Ivliev, Hendrik G. Stunnenberg, Mattia Frontini, Mikhail Spivakov, Peter Fraser, Antony Cutler, Chris Wallace, Stephan Beck

## Abstract

Understanding the regulatory landscape of human cells requires the integration of genomic and epigenomic maps, capturing combinatorial levels of cell type-specific and invariant activity states.

Here, we segmented whole-genome bisulfite sequencing-derived methylomes into consecutive blocks of co-methylation (COMETs) to obtain spatial variation patterns of DNA methylation (DNAm oscillations) integrated with histone modifications and promoter-enhancer interactions derived from promoter capture Hi-C (PCHi-C) sequencing of the same purified blood cells.

Mapping DNAm oscillations onto regulatory genome annotation revealed that enhancers are enriched for DNAm hyper-oscillations (>30-fold), where multiple machine learning models support DNAm as predictive of enhancer location. Based on this analysis, we report overall predictive power of 99% for DNAm oscillations, 77.3% for DNaseI, 41% for CGIs, 20% for UMRs and 0% for LMRs, demonstrating the power of DNAm oscillations over other methods for enhancer prediction. Methylomes of activated and non-activated CD4^+^ T cells indicate that DNAm oscillations exist in both states irrespective of activation; hence they can be used to determine the location of latent enhancers.

Our approach advances the identification of tissue-specific regulatory elements and outperforms previous approaches defining enhancer classes based on DNA methylation.

## Background

Enhancers are regulatory DNA elements that can increase the transcription of genes under their control in a cell type-specific manner in conjunction with transcription factors, chromatin states and a variety of cofactors. They were discovered over 30 years ago in simian virus 40 and today account for the majority of regulatory sequences in mammalian genomes [1,2]. There are multiple classes of enhancers [1] and new ones are still being identified, including the recently described latent [3] and super [4] enhancers as well as a class of as yet unknown function defined by acetylation of histone globular domains (H3K64ac and H3K122ac) rather than histone tails [5]. Because of their fundamental role in gene regulation, one of the key questions has been how to identify all enhancers using either computational or experimental approaches [6]. Towards this aim, genome-wide approaches have been developed based on DNA sequence conservation, profiling of histone modifications, transcription factors, transcription coactivators, chromatin states, enhancer RNAs (eRNA) and chromosome conformation capture [reviewed by 7].

DNA methylation is the most ubiquitous and stable epigenetic mark and is known to exert both positive and negative control over gene expression [8]. Here, we assessed the predictive potential of DNA methylation marks for identifying enhancers. We leveraged 24 whole-genome bisulfite (WGBS) methylomes generated by the BLUEPRINT Project for 9 purified haematopoietic cell types, including activated and non-activated CD4^+^ T cells, to address three fundamental questions:Can enhancers be predicted based on DNA methylation data only? [ii] How do the results compare to those obtained with traditional predictions using histone modifications (H3K4me1, H3K4me3, H3K27ac, H3K36me3), chromatin states (ChromHMM) and more recent methods that analyse chromosomal interactions between promoters and enhancers such as PCHi-C [9–12] ? [iii] Based on comparative methylome analysis before and after cell activation [13], can latent enhancers be predicted prior to their activation? In the context of the immune cells analysed here, the latter would for instance allow assessment of how well in vitro activation mimics an in vivo immune response.

In on our previous work [14] we found that certain patterns of DNA methylation could be used to infer the location of enhancers. By segmenting methylomes into consecutive blocks of comethylation (COMETs), we observed spatial oscillatory methylation signatures that associated with regulatory regions. These oscillatory signatures, termed harmonics or DNAm oscillations can be estimated using the the COMETgazer algorithm [14] as a continuous CpG density-independent K-period percentage difference series based on the continuous smoothed methylation level estimate. While sudden breaks in oscillation patterns can be used to define COMETs, in this study we used high levels of oscillations (hyperoscillatory patterns) to analyse methylomes specifically for enhancers in the context of the questions posed above.

## Methods

### Datasets

The BLUEPRINT DNA methylation and ChIPSeq data EGA numbers are summarized in **Supplementary Table 1**. Raw sequencing reads for PCHi-C data have been submitted to EGA (http://www.ebi.ac.uk/ega), accession number EGAS00001001911. DNA methylation and ChIPseq data for the CD4^+^ T cell activation are available as raw sequencing reads (submission to EGA in progress).

### Genome segmentations

The 2014 Blueprint segmentations were generated using ChromHMM version 1.12 (http://compbio.mit.edu/ChromHMM/) by the Centro Nacional de Investigaciones Oncológicas (CNIO). They were only run on samples with full reference epigenome histone modification set and matched input on GRCh37 (http://dcc.blueprint-epigenome.eu/&#/md/secondary_analysis/Segmentation_of_ChIP-Seq_data_20150128).

The Ensembl Regulation team has run the Ensembl Regulatory Build on the 2015 BLUEPRINT segmentations merging in data from ENCODE segmentations, DNase and CTCF peaks [15] (http://dcc.blueprint-epigenome.eu/&#/md/secondary_analysis/Ensembl_Regulatory_Build_20150128). The Blueprint segmentation pipeline using ChromHMM requires the existence of all the individual specific reference epigenome ChIP-seq histone modifications (H3K3me1, H3K4me3, H3K9me3, H3K27ac, H3K27me3, H3K36me3) and Input (Control). The Regulatory Build was run using the resulting segmentations.

The GRCh37 segmentations are available at: ftp://ftp.ebi.ac.uk/pub/databases/blueprint/releases/20150128/homo_sapiens/secondary_analysis/Segmentation_of_ChIP-Seq_data/ and the builds at:ftp://ftp.ebi.ac.uk/pub/databases/blueprint/releases/20150128/homo_sapiens/secondary_analysis/Ensembl_Regulatory_Build/.

For the PCHi-C analysis, we ran 2015 Blueprint segmentations with additional states, as we could not identify any poised states from the CNIO segmentations, and we ran the Regulatory Build again using these segmentations. The segmentation data are available from: ftp://ftp.ebi.ac.uk/pub/contrib/pchic/Blueprint/.

### BLUEPRINT DNA methylation pipeline

The BLUPRINT methylation pipeline is available at:http://ftp.ebi.ac.uk/pub/databases/blueprint/releases/20140811/homo_sapiens/README_bisulphite_analysis_CNAG_20140811.

LMR and UMR were defined using the Bioconductor tool methylSeekR (https://bioconductor.org/packages/release/bioc/html/MethylSeekR.html). DNAase clusters from 125 cell types was obtained from UCSC at http://hgdownload.cse.ucsc.edu/goldenPath/hg19/encodeDCC/wgEncodeRegDnaseClustered/ including DNase clusters (V3) from UW and Duke ENCODE data uniformly processed by the ENCODE Analysis Working Group.

### Harmonic analysis

For the exploratory analysis (ROADMAP and BLUEPRINT segmentations), fixed genome windows of 1000 bp and 10% oscillations in DNA methylation were used. For ROADMAP analyses, we defined enhancers as PCHi-C regions overlapping with histone mark signal (+/− 500 bp) with the following definitions: active (strong H3K4me1, strong H3K27ac), poised enhancer (strong H3K4me1 and weak H3K27ac), promoter-proximal (near region with strong H3K4me3, H3K27ac and depleted in H3K4me1), transcribed (strong H3K36me3) and quiescent (no noticeable active histone marks). The quiescent regions were used for background, in order to calculate the enrichment ration of oscillation counts.

DNAm oscillations (harmonics) were defined as a continuous CpG density-independent K-period percentage difference series based on the continuous smoothed methylation level estimate. The quantile distribution of DNAm methylation values is analysed independently for each chromosome. Most of the oscillations are around zero, and these define regions of co-methylation. Fragmentation in the methylome structure is defined as significant deviations in the quantile distribution used to call individual regions of co-methylation. This is as described in Libertini et al. [16](http://www.nature.com/articles/ncomms11306). By counting the number of significant DNAm oscillations, the total number of counts was defined as harmonics count per genomic window. Throughout the manuscript, harmonics refer to the number of DNAm oscillation counts per region of interest. The analysis used 2015 BLUEPRINT segmentations (http://www.ensembl.org/info/genome/funcgen/regulatorysegmentation.html) and the ENSEMBL regulatory build (http://www.ensembl.org/info/genome/funcgen/regulatory_build.html). This was run using fixed windows of 3000bp with oscillations of at least 1% to define harmonics [16]. HMM_state, PCHi-C and histone_mark regions were defined by overlap, for this region some windows are transitions and are assigned to more than one category. Regions of high, medium and low harmonics were defined using > 10 (high) and < 3 (low) harmonics per genomic window as a hard threshold for all samples. The medium harmonics are between 3 and 10. Regions of highest oscillatory signal (hyperoscillatory regions) were defined as the 99th quantile of the harmonics distribution in each sample. Putative regions were defined as devoid of any PCHi-C or histone mark signal with the exception of H3K36me3, in order to capture transcribed regions which may also have a regulatory function.

### BLUEPRINT ChIP-Seq analysis pipeline

The details of the pipeline are available at: http://dcc.blueprint-epigenome.eu/#/md/chip_seq_grch37.

### ChIP-Seq analysis (CD4^+^ T cells)

ChIP-seq reads for histone modifications and control assays were mapped to the reference genome using BWA-MEM (https://arxiv.org/abs/1303.3997). Samtools (http://bioinformatics.oxfordjournals.org/content/25/16/2078.short) was employed to remove secondary and low-quality alignments (PHRED score <= 40 or no bits matching the SAM octal flag 3 or some bits matching the octal flag 3840). Remaining alignments were sorted and indexed.macs2 (https://genomebiology.biomedcentral.com/articles/10.1186/gb-2008-9-9-r137) was used to call peaks on each histone modification sample. Its broad peak mode was used for H3K27me3, H3K36me3, H3K9me3 and H3K4me1. In all cases, the fragment size parameter was estimated with PhantomPeakQualTools (http://www.g3journal.org/content/4/2/209.long). Differential peaks for each paired non-activated vs activated histone mark on CD4 T cells were obtained with THOR (http://nar.oxfordjournals.org/content/early/2016/08/01/nar.gkw680.full).

### PCHi-C analysis pipeline

Interaction confidence scores were computed using the CHiCAGO pipeline (Cairns et al., 2016). Briefly, CHiCAGO calls interactions based on a convolution background model reflecting both ‘Brownian’ (real, but expected interactions) and ‘technical’ (assay and sequencing artefacts) components. The resulting p-values are adjusted using a weighted false discovery control procedure that specifically accommodates the fact that increasingly larger numbers of tests are performed at regions where progressively smaller numbers of interactions are expected. The weights were learned based on the decrease of the reproducibility of interaction calls between the individual replicates of macrophage samples with distance. Interaction scores are then computed for each fragment pair as –log-transformed, soft-thresholded, weighted p-values. Interactions with a CHiCAGO score ≥ 5 in at least one cell type were used for further analysis as in Javierre et al (in press). Raw sequencing reads have been submitted to EGA (http://www.ebi.ac.uk/ega), accession number EGAS00001001911. High-confidence interactions (CHiCAGO score >= 5 in at least one cell type) are available via the CHiCP browser, where they can be visualised alongside GWAS data (http://www.chicp.org) and as custom tracks for the Ensembl browser (ftp://ftp.ebi.ac.uk/pub/contrib/pchic/CHiCAGO). The regulatory build annotations and segmentations of the BLUEPRINT datasets are available as a track hub for the Ensembl browser (ftp://ftp.ebi.ac.uk/pub/contrib/pchic/hub.txt). Further processed datasets, including TAD definitions, regulatory region annotations, specificity scores and gene prioritization data, are available via Open Science Framework (https://osf.io/u8tzp).

### Machine learning

Partial Least Squares (PLS), Generalized Linear Models (GLM), and Elastic Net (SN) machine learning methods incorporating a cross-validation framework were build in order to assess the predictive power of features including PCHi-C targets, four histone marks and DNAm oscillations. Although more generally labelled as ‘regression’, the approaches used here may be naturally extended to classification problems. The three approaches here differ in the way they capture model parameters and feature importance. For PLS, features are combined into latent variables. GLM uses stepwise addition/removal of features to identify what is considered to be an optimal subset. EN performs shrinkage and variable selection to improve predictive performance and interpretation of the model.

Performance was assessed using area under the ROC curve (AUC) computed by a cross-validation procedure in which features to be selected were taken into account. Logistic regression was used for forward selection in order to rank the features. Elastic net regression was also run and features were ranked based on the final model coefficients.

### Partial least squares regression

A machine learning model was built using partial least squares (PLS) regression [17]. PLS regression is a powerful method for building predictive models with many factors that are highly collinear and functions by constructing a multivariate linear regression model by projecting a set of predictor variables and a response variable onto a new space. This constructs a matrix of latent components that are linear transformations of the original predictor variables and maximize the explained multidimensional variance direction in the response variable space. These latent variables are then used for prediction in place of the original variables [18]. In our scenario in which the response variable was categorical, a variant of PLS regression termed PLS-DA (discriminant analysis) was used. The predict function from the caret package was used to train PLS classifiers.

### Generalized Linear Models

Forward stepwise regression using a generalized linear model (GLM) [19] was used to model the relationship between the predictor and response variables. The GLM framework provides for Normal, Binomial, Poisson and Multinomial likelihoods, with a variety of link functions. Here, since the response variable was categorical, a binomial model and the logit link function was used (logistic regression). The model started with no variables and the effect of the addition of each variable was estimated at each step. Variables were added to the model according to their ranked t-statistic until no further improvement was seen. Here, performance was assessed using the AUC

### Elastic Net

Elastic net (EN) [20] was additionally used to build predictive models in a classification setting. Least-squares estimates of regression coefficients may be highly unstable, especially in cases of correlated predictor variables, leading to low prediction accuracy. Shrinkage methods (setting some of the regression coefficients to zero) e.g. lasso regression [21] may result in estimates with smaller variance and improved accuracy. Additionally, EN facilitates variable selection by encouraging a sparse solution and thus retaining only important predictor variables in the model. In addition to a sparse solution, EN can encourage group selection amongst correlated variables (which may be considered an advantage in a biological setting). EN requires parameters alpha and lambda to be defined. Alpha sets the degree to which the penalization is more towards the L2 (ridge regression) and L1 (lasso regression). Lambda is the shrinkage parameter and controls how strongly the coefficients are shrunk (where 0 would result in no shrinkage). In this study both alpha and lambda were optimized through an inner CV loop where for each of the CV runs performed, an additional 5 fold CV was performed to tune these parameters using a grid search.

### Cross Validation

To enable for an accurate estimate of model performance, the analysis was performed using a balanced (stratified) nested cross-validation where the proportion of positive and negative controls was maintained during each iteration. In the inner loop of the cross-validation, we performed feature selection using recursive feature elimination approach. In the outer loop, we tested performance of the resulting classifiers after training them on the best performing feature set from the inner loop. Both the outer and the inner loops represented 5-fold cross-validations.

Negative controls were defined as a random subset of all genome locations that were not positive controls (enhancers). This allowed for avoiding a strong imbalance between the number of positive and negative controls during the learning process. The negative controls were defined in each fold of the outer loop independently, so the effect of random selection was averaged out across the cross-validation. The number of negative controls was set to 10 times the number of positive controls.

The entire cross-validation – from splitting samples in the outer folds up to the integrative scoring – was repeated 10 times. PLS scores were converted into genome-wide ranks within each repeat. The ranks were finally averaged across the repeats to produce a final region ranking.

### Recursive feature elimination

Feature selection was performed within the inner loop of the cross-validation as follows. For a given outer cross-validation fold:

- execute an inner cross-validation based on 80% of samples and the complete feature set:

– Determine an optimal number of PLS components (ncomp)
– For the model with the optimal ncomp, store feature weights and PLS performance
- remove 10% features with lowest weights
- re-run the inner cross-validation using remaining features

– Determine an optimal ncomp
– For the model with the optimal ncomp, store feature weights and PLS performance
- repeat until only one feature remains
- determine the feature set at which best performance was achieved during the cross validation

We then applied the model with best performance to the left-out samples from the outer loop to measure the model performance. Feature selection was performed 50 times: 10 repeats multiplied by 5 outer folds. For each feature, we tracked the number of times it was selected in the final model (a value from 0 to 50). This measure reflected feature importance during the modeling process.

### Estimation of variability in DNAm oscillations between cell types

Using the same count data which was used to generate Figures 2 and 3 (i.e. distribution of DNAm oscillation counts across the genome at a 3000 bp resolution), we used a negative binomial model to estimate consistent differences between samples using replicates (Supplementary Table 1) using DESeq2 [22] with adjusted p-value < 0.1 (BH) and absolute logFC > 0. The software estimates low counts and outliers, which may influence the results.

### Enrichment analysis

Enrichment analysis methods have become commonplace tools applied to the analysis and interpretation of biological data. The goal is to discover pathways or processes associated with the gene list of interest. The significance of enrichment is defined by using the hypergeometric test [23], using the following parameters:

N – number of network objects covered by the whole ontology
R – number of network objects in a list under analysis
n – number of network objects associated with a particular category from the ontology
r – number of gene from input list intersecting with genes from a particular category

As a result, all terms from the ontology are ranked according to calculated p-values. Ontology terms with p-values less than the p-value threshold 0.05 are defined as statistically significant and therefore relevant to the studied list of genes. In other words, the gene list is associated with a quantitatively ranked list of pathways and processes summarizing its effects at a systems-biology level.

For this part of the project, we used Pathway Maps ontologies (maps) along with process networks, and three levels of canonical gene ontologies (biological process, molecular functionsm, cell localizations) Clarivate analytics canonical pathway maps represent images of signaling pathways describing a particular biological mechanism. Pathway maps comprehensively cover human, mouse and rat signaling and metabolism.

Annotations were based on the Clarivate Analytics priorietory database Metabase which underlies the integrated software suite MetaCore for functional analysis of Next Generation Sequencing, gene expression, CNV, metabolic, proteomics, microRNA, and screening data. MetaCore is based on a high-quality, manually-curated database of molecular interactions, molecular pathways, gene-disease associations, chemical metabolism and toxicity information (http://clarivate.com/?product=metacore).

Metabase relations between molecular entities: interactions, associations, reactions An interaction in MetaBase describes an influence an object has on another object. There are a few main types of interactions between molecular entities:

- Protein-protein interactions. This type of interaction constitutes the majority of signaling networks
- RNA – protein interactions. A variety of interactions mostly describing protein translation, RNA degradation, and interactions of miRNA with its protein targets
- RNA – RNA interactions. Interactions between different types of RNA (for instance, in the ribosome). Also includes micro RNA – target mRNA interactions
- Compound – protein interactions. Interactions of small molecular endogenous and xenobiotic ligands with proteins leading to modulation of proteins activity
- Compound – DNA, RNA interactions. Mostly unspecific interactions of planar or highly reactive compounds leading to interruptions in gene expression
- Compound-compound interactions. These are indirect interactions between bioactive compounds: drug-drug interactions and endogenous ligand – synthetic ligand interactions

In MetaBase, all interactions are attributed with a 1) direction, indicating signal transduction, 2) effect, depicting character of influence (e.g. inhibition, activation), 3) mechanism, showing how the effect has been reached, 4) experimental details from literature source, confirming the interaction, and 5) trust, given by an expert and indicating the probability of the interaction’s existence. Based on the mechanism, interactions are divided on direct interactions meaning that physical contact between interacting objects occurs and indirect interactions, when observing effect between the objects is mediated by omitted interactions or a whole pathway.

In MetaBase, the vast majority of interactions are directional, i.e. depict “from – to” relations. This is characteristic for individual interactions such as microRNA – target inhibition, as well as interactions linked into multi-step linear signaling or metabolic pathways. It initiates with a ligand – receptor interaction on the cellular membrane and is transmitted via several signal transduction interactions to the transcription factor, followed by its binding to the promoter of a target “effector” gene such as endogenous metabolic enzymes. Information on the direction of interaction is not always available from experimental literature. For instance, yeast-2-hybrid or co-immunoprecipitation assays can only establish the fact of binding but not the direction of interaction. In such cases, the direction can be established by its relative position in the signaling pathway. When no additional data on the pathways is available (typical case for recently annotated proteins), the direction is not marked until more data establishes it.

### Segmentation with MethylSeekR

Unmethylated and low methylated regions were estimated with *MethylSeekR* [24] with 5% FDR cutoff and m=0.5 as input parameters.

## Results

The data used in this study comprised matched samples of whole-genome bisulfite sequencing (WGBS) and histone mark ChIP-seq (H3K4me1, H3K4me3, H3K27ac, H3K36me3) for 24 healthy donors and target regions (promoter interacting) from promoter capture Hi-C (PCHi-C) for 9 primary haematopoietic cell types, including neutrophils, monocytes, CD4^+^ naïve T cells, CD8^+^ naïve T cells, erythroblasts, megakaryocytes and macrophages (**Supplementary Table 1**) (**Methods**). We also analyzed methylomes of activated and non activated CD4^+^ T cells from Burren et al. [13]. Using these datasets, we tested the hypothesis that patterns in the spatial oscillations of DNA methylation (DNAm oscillations) may be used to predict enhancers discriminate them from different types of the genomic regions, showing that enhancers display a hyperoscillatory pattern. In order to establish the predictive power of DNA methylation, we compared the methylome oscillatory signature (harmonics) [14] using three different genome segmentations, which were used as reference sets. One set was based on criteria established by Roadmap Epigenomics Project [25] (**Methods**), and two were internal BLUEPRINT analyses (**Methods**).

### Enhancers are enriched in DNAm oscillations

Initially, we used enhancer definitions established by the Roadmap epigenomics project [25,26]. Using these criteria, we defined a benchmark set of different region types using histone marks and PCHi-C data (**Methods**). Using this set, we investigated the predictive nature of DNA methylation by defining genomic classes based on the frequency of DNAm oscillations *(harmonics)*.

Across all 24 samples, we found the genome coverage of enhancers to be on average 4% [26], with 1% of regions covering promoter proximal, 15% to be transcribed regions, 66% defined as quiescent regions and 24% of the genome being devoid of signal. Given these genomic segmentations, we found striking differences in the pattern of DNAm oscillation differences between them. Indeed, on average 30% of harmonics were in enhancer regions, 10% in promoter proximal, 45% in transcribed regions and 15% in quiescent regions, though their frequency (i.e. the number of harmonics per window) varied across genomic region types (**Supplementary Table 2**). We found that different region types had distinct DNAm oscillation profiles (**Supplementary Table 2**). Based on this, we speculated that the frequency of oscillations is informative and it may be a predictor of region type, a hypothesis which we later tested using various machine learning algorithms (see **Methods** and *machine learning* section).

Using Roadmap enhancer regions definitions, we found enrichment (>30 fold) of harmonics (hypergeometric test, P < 0.001), with oscillations of magnitude of at least 10% in DNA methylation levels as defined in [14] within enhancer regions across all cell types, where quiescent regions had very little signal (**Supplementary Table 2**). Here, the different types of regions had distinct oscillatory profiles, where active enhancers and promoter proximal regions showed the highest number of harmonics in every cell type, while transcribed regions and quiescent regions showed the lowest number of harmonics (**Supplementary Table 2**). Interestingly, poised enhancers showed fewer harmonics than active enhancers, but on average twice as many as transcribed regions (**Supplementary Table 2**).

As a second analysis, we investigated the harmonics content of methylomes by using genomic region definitions based on the BLUEPRINT genome segmentations (**Methods**). These segmentations were run using the collective set of BLUEPRINT data with the ChromHMM software with the purpose of partitioning the genome based on predicted genomic region function. This dataset included a predicted range of different enhancer classes that we used as benchmark for our analysis (**Methods**). As in the first analysis, we found that the harmonics signature of 10% oscillations could be used to discriminate different classes of enhancers (**Supplementary Figure 1**). Here, each of the described categories displayed a distinct distribution of harmonics. This test set validated our initial observations of hyperoscillatory patterns in enhancers. For this reason, we proceeded to test whether harmonics could be employed as a predictive tool for enhancer characterization using several other datasets to support this model.

### Predicting genomic regions using DNAm oscillations in comparison to genome segmentations, PCHi-C and histone marks

The focus of our third analysis was based on the BLUEPRINT segmentations integrated in conjunction with the ENSEMBL Regulatory Build [27] with additional states with respect to the previous analysis shown in **Supplementary Figure 1**. Here we used DNAm oscillations at a higher level of resolution, with differences of 1% being used to define harmonics (**Methods**).

In this setting, genome segmentations were taken from the recently published ENSEMBL Regulatory Build [27,28], which comprises publicly available data from different large epigenomic consortia (including ENCODE, Roadmap Epigenomics and BLUEPRINT) and includes BLUEPRINT chromatin segmentations (http://www.ensembl.org/info/genome/funcgen/regulatory_segmentation.html) The ENSEMBL Regulatory Build includes 8 chromatin states, including **heterochromatin**, **repressed**, **gene** *(genic)*, **weak** *(weak enhancers)*, **distal** *(distal enhancers of moderate signal)*, **proximal** (**proximal enhancers of moderate signal**), **poised** (which are in fact *enhancers of strong active signal)* and **tss** *(transcription start sites)*. The regions defined as **poised** in this classification include H3K27ac signal (i.e. active enhancers). For simplicity, we have retained the same nomenclature for the labels used to functionally annotate the states as designated in the Regulatory Build (http://www.ensembl.org/info/genome/funcgen/regulatorybuild.html) [27, 28].

We investigated the harmonics of each of the region types based on three reference sets, which in turn were based on three different types of overlaps. At the most basic level (first reference set), region definitions were based on the genome segmentations (*regions α*). As a second reference set, we took the overlap of these with regions where PCHi-C signal [28] existed for targets (enhancers; *regions β*) (**Methods**). As a third reference set, we took a further subset of regions where both PCHi-C and specific patterns of histone marks signal existed (*regions γ*) (see PCHi-C section in **Methods**). Based on these three reference sets, we expected regions α to have the least confidence for predicting enhancers, because they were only based on a computational model. Conversely, we expected the regions γ to have the strongest confidence because they were based on the overlap of all reference datasets.

Indeed, across over all 24 samples the overall enrichment of harmonics over the background (heterochromatin regions) improved when comparing the three reference sets (**Figure 1A**). This is particularly true for enhancers of strong signal (termed **poised** by Cunningham et al. [27]) and tss. When comparing the three reference sets, each genomic region type has distinct harmonics, enrichment and genome coverage (**Supplementary Figures 2-3**). In general, enhancer regions show low coverage and high enrichment of harmonics while heterochromatin regions show the opposite pattern (**Supplementary Figures 2-3**). The three reference sets show comparable levels of harmonic content, with the exception of **poised** regions, where using three sets of overlap (regions α, β and γ) greatly increases the ratio of harmonics enrichment (**Figure 1A**). Taken together, these results show that high frequency oscillations are associated with different enhancer and promoter proximal classes, even as these have lower genome coverage (**Supplementary Figure 2**).

**Figure 1.**
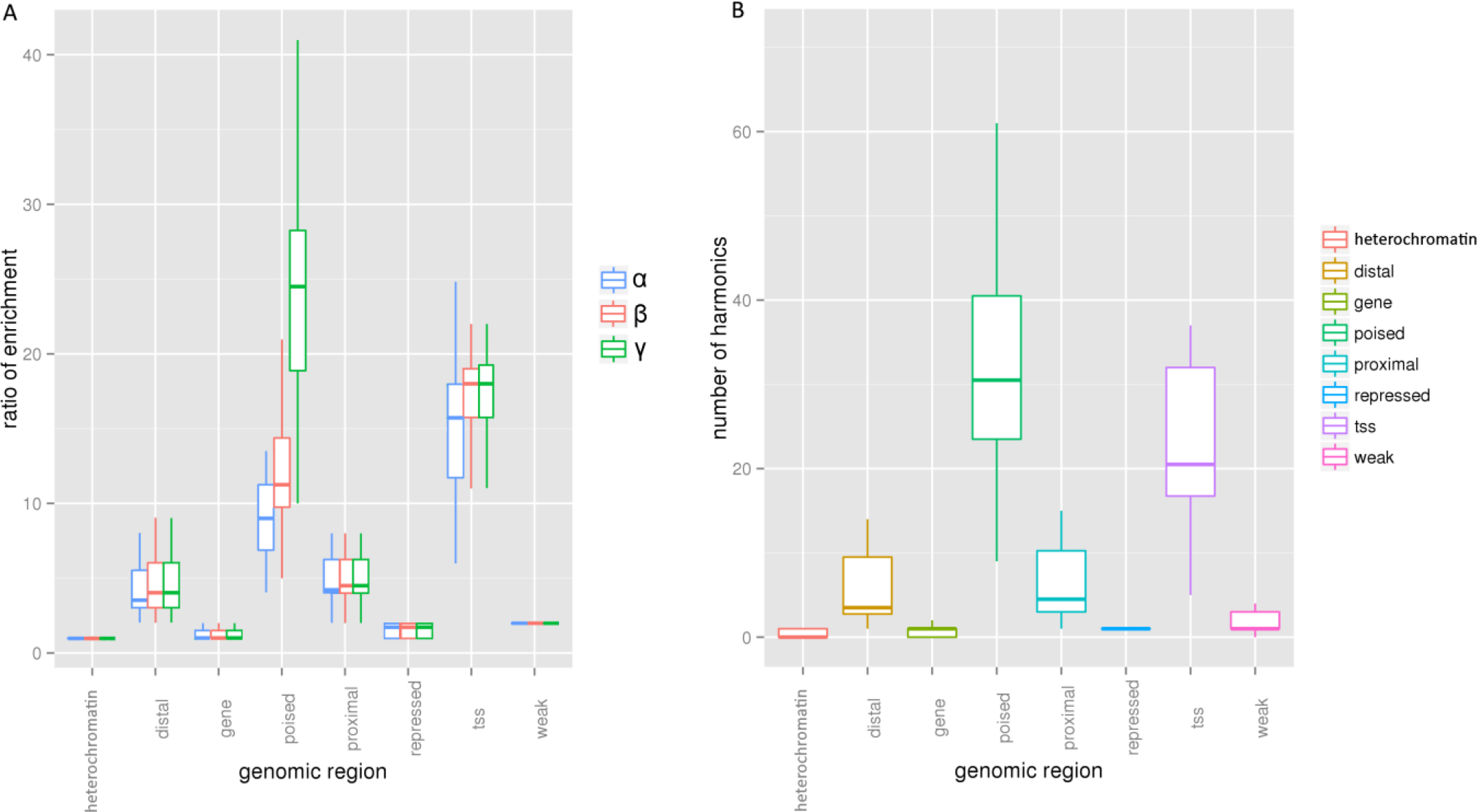
(A) Enrichment of harmonics over the background (heterochromatin regions) based on BLUEPRINT segmentations integrated in conjunction with the ENSEMBL Regulatory Build [27]. Region definitions are based on the genome segmentations (*regions α*), overlap of these with regions where PCHi-C signal existed for targets (*regions β*), a further subset of regions where both PCHi-C and histone mark signal exists (*regions γ*). (B) Number of harmonics in high confidence regions based on histone marks using BLUEPRINT segmentations integrated in conjunction with the ENSEMBL Regulatory Build [27] (Methods).

Across all samples, the genome-wide Pearson correlation between harmonics frequency and the position of the **poised** class is 0.4 (p < 0.001), with **tss** 0.5 (p < 0.001), **distal** 0.25 (p < 0.001), **proximal** 0.1 (p < 0.001), **heterochromatin** −0.2 (p < 0.001), **repressed** 0 (p < 0.001) and **gene** 0 (p < 0.001). By randomizing the class labels as a negative control, this association is lost (p > 0.05).

The high confidence reference set (*regions γ*) showed a clear discrimination in the harmonics pattern (**Figure 1B**), supporting our hypothesis that harmonics content can be used to discriminate different classes of genomic regions, and that enhancers show elevated levels of DNAm oscillations. This pattern is particularly evident when assessing quantile distributions of harmonics content, where the differences in distribution were associated to different enhancer types. This is a striking pattern which seems to be reproducible in each cell type analysed (**Figure 2A**). This phenomenon is also reproducible across all samples, as evident from an overall analysis (**Figure 2B**).

**Figure 2.**
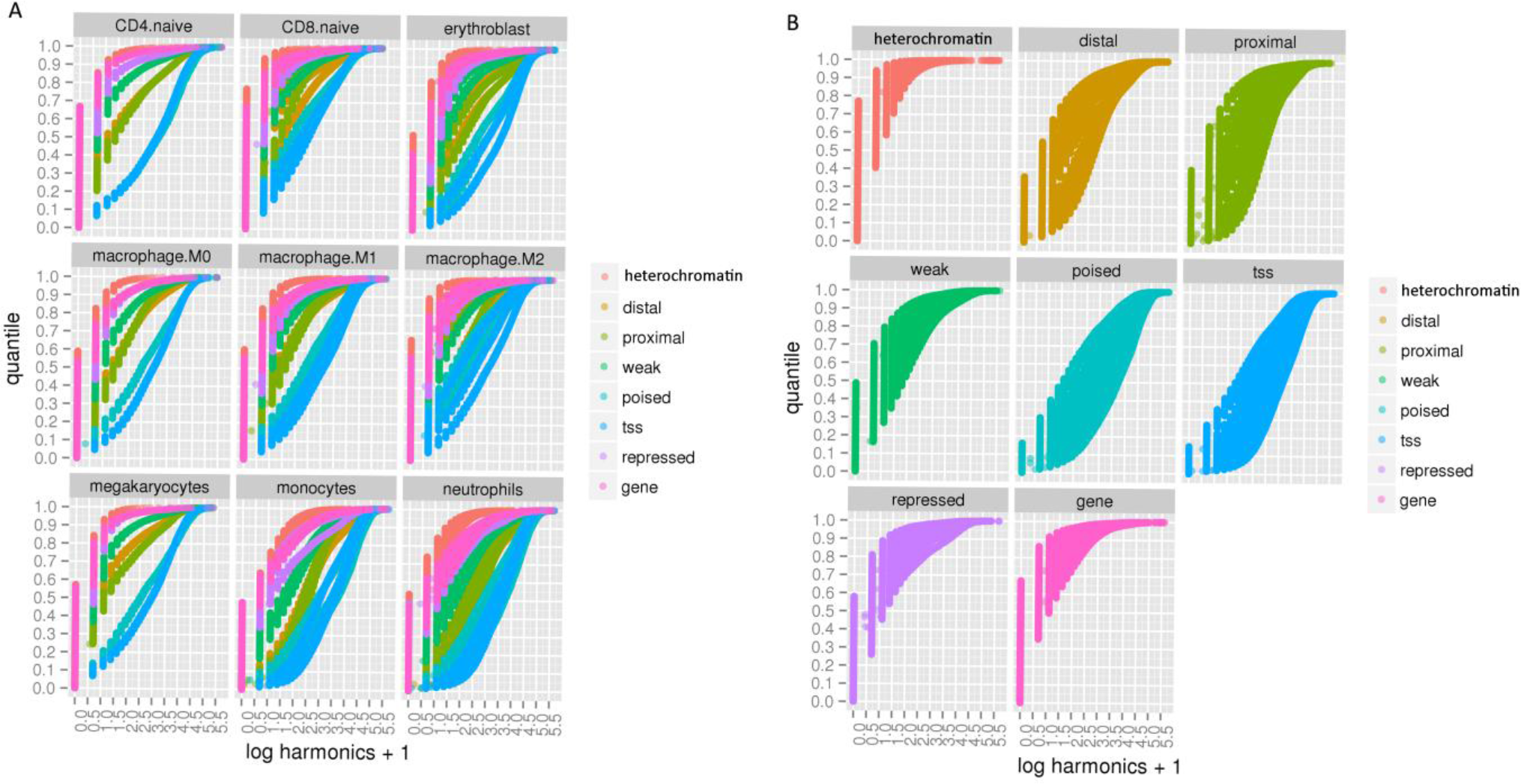
(A) Quantile plots of harmonics distributions showing distinguishable patterns in harmonics content, where each region is defined by a different incline and this pattern is reproduced in each cell type. (B) Quantile plots of harmonics distributions: each type of region is defined by a different incline, specific to oscillatory content, where regulatory regions are enriched in harmonics.

Using these quantile distributions in order to quantify the oscillatory effect, we calculated the differences between the integrals. Across all the cell types, the integrals of the areas under the curve are distinct for each region type. Our data show a larger effect for the **poised** (~34.9) and **tss** (~35.7), while **distal** is the second largest (~15.1), then **weak** (~7.8), **repressed** (~5.5), **gene** (~3.4) and **heterochromatin** (~1.4). With this numerical scale, it was possible to quantify and rank the oscillatory signature of each of the enhancer classes and associate it with an area size. According to this analysis, enhancer and tss regions show a harmonics signature (i.e. integral) which is 30-fold greater than the background, supporting our earlier claim (**Figure 2B**).

### Strong recovery of enhancers using DNAm oscillations only

Next, we investigated how many of the high confidence regions *(histone_marks)* could be recovered with DNAm oscillations alone. We found that using high and medium harmonic signal (**Methods**) we could on average recover 90% of **poised**, 85% of **tss**, 57% of **distal**, and 60% of **proximal** enhancer regions (**Figure 3**). The oscillatory signature of **weak** enhancer regions was too subtle to be effectively used, even if it was statistically different (Mann-Whitney U, P< 0.001) from **heterochromatin**, **repressed** and **gene** regions (**Figure 1B**). The weak enhancer regions were on average too many for the signal to be strong enough (**Supplementary Figure 2**).

**Figure 3.**
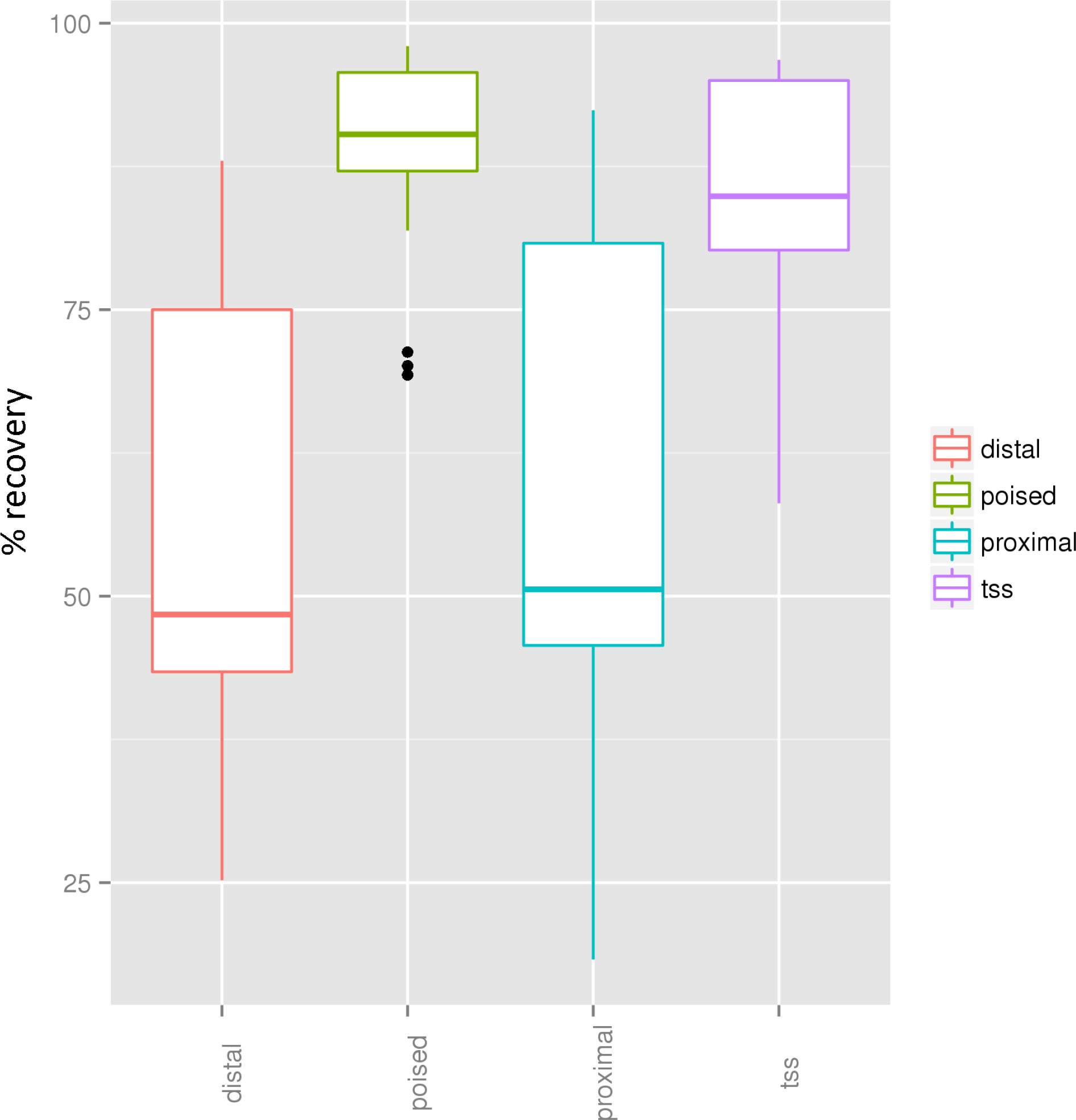
Recovery of enhancers using DNAm oscillations alone. We found that using high and medium harmonic signal, we could on average recover 90% of poised, 85% of tss, 57% of distal, and 60% of proximal enhancer regions. The data is represented with boxplots as mean +/− 95% confidence interval.

We found some variation in recovery among different cell types, depending on the base oscillatory signature of the background, where neutrophils and monocytes had the strongest signal, and CD8+ naive T cells and megakaryocytes the weakest (**Supplementary Figure 4**), with the lowest number of high harmonic counts and 0 oscillations in the background (**Supplementary Table 3**). It is interesting to note that variability between cell types showed DNAm oscillations to have the greatest difference between neutrophils and monocytes. In this regard, we estimate the effects of variability of DNAm oscillations between cell types at individual loci (**Methods**). The analysis represents consistent variability among cell types using biological replicates (Supplementary Table 1). Using the same count data which was used to generate Figures 2 and 3 (i.e. distribution of DNAm oscillation counts across the genome at a 3000 bp resolution), we established that within the lymphoid lineage, DNAm oscillations predicting enhancers varied 11% between CD4^+^ naive T cells and cytotoxic CD8+ T naive cells. Within the myeloid lineage, DNAm oscillations megakaryocytes and erythroblasts varied 27%. Subtle differences between resting (M0), activated (M1) and alternatively activated (M2) macrophages were also reported at ~0.03% variability in DNAm oscillations, showing the high resolution potential of this method. The full results of this analysis are shown in supplementary Table 9.

### Putative enhancers defined by methylation alone

Regions of high harmonics (> 99th quantile) with no identifiable histone marks or PCHi-C signal were hypothesized to carry enhancer signal, and these were quantified for each of the samples. On average 1% of hyperoscillatory regions was not assigned to any known enhancer category based on chromatin segmentation (**Supplementary Table 1**), amounting to a sum of 1630 putative enhancers across cell types. In addition, we found 238 regions devoid of histone marks, not assigned to any category, and with a hyperoscillatory signature which overlapped with the targets (enhancers in PCHi-C promoter enhancers interactions) (**Supplementary Table 1**). Using the PCHi-C data, we found the associated genes by overlap of +/−2500 bp of the associated promoter. While 62% of genes where found to be inside the regions thus defined, 10% of genes where found to be downstream, 18% to partially overlap and 10% to be upstream of these regions, where the median distance from the TSS was 10,000 bp. In absence of any gene expression data, and in order to ascribe a biological signature to these regions by categorizing genes into groups, we ran functional overview analyses using *Clarivate Analytics* tools for pathway analysis and gene ontology enrichments (http://clarivate.com/life-sciences/discovery-and-preclinical-research/metacore/). By doing so, we identified enriched pathways associated to this global signature across cell types (**Supplementary Table 4**). In addition, we ran a cell specific analysis for macrophages which had the largest number of samples (**Supplementary Table 1**). Interestingly, these analyses yielded significant results for enriched pathways. Results included pathways specific to the immune response such as IL-4 signalling (**Supplementary Figure 5**), as well pathways specific to blood cell types (**Supplementary Figure 6-8**). While gene ontology analyses confirmed that the most significant biological processes are associated with transcription (**Supplementary Table 5**), the genes associated with this signature counted enrichment for protein kinases and microRNAs. Among these, microRNAs involved in epithelial-to-mesenchymal transition were identified as bearing hyperoscillatory signatures in their promoter regions which are devoid of histone marks (**Supplementary Figure 6**). In addition to these, one of the most significantly enriched pathways relates the immune response in macrophages, where several of the kinases bear this epigenetic signature (**Figure 4**). Putative enhancers from macrophage samples were associated to genes in macrophage-specific pathways (**Figure 4**), showing a degree of cell-type specificity.

**Figure 4.**
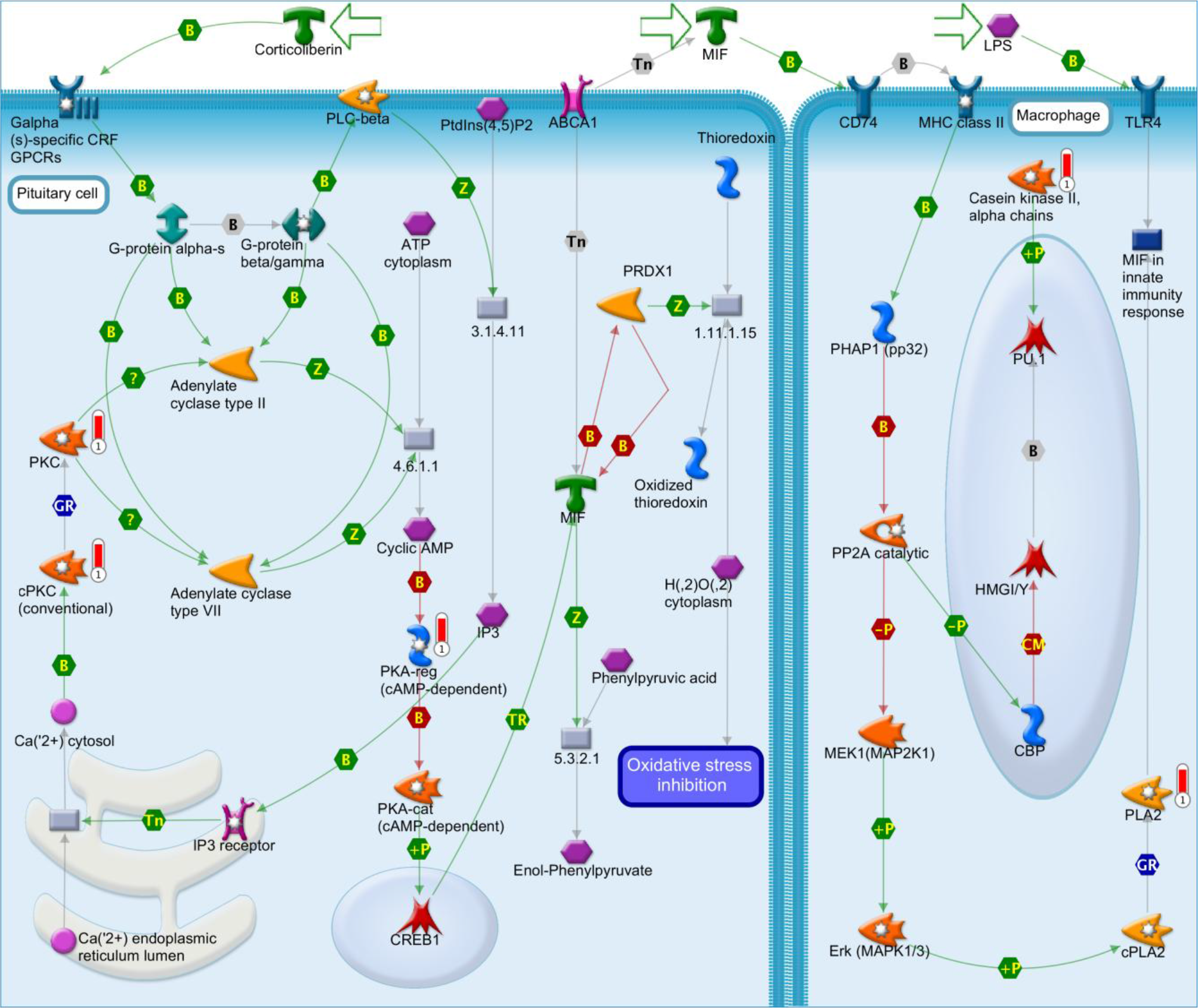
Immune response_MIF - the neuroendocrine-macrophage connector pathway. Genes highlighted with red bar are associated with putative enhancers identified with DNAm oscillations.

### CD4 activation patterns can be recapitulated by DNA methylation

We explored the DNAm oscillatory signature of activated and non activated CD4^+^ T cells, which, as already discussed, bear strong transcriptional and epigenetic differences [13]. DNAm oscillations in CD4^+^ T cells recovered ~90% of enhancer regions in both non-activated cells and activated cells, when using previously established histone mark definitions as reference set and 1% DNAm oscillations and 3000bp windows (**Table 1**).

**Table 1.** **Prediction of CD4^+^ T cell activation enhancers** based on DNAm oscillations, as validated by enhancer definitions based on corresponding histone mark data. **NAC_histones**: enhancer regions as defined by histone marks only (non-activated CD4^+^ T cells).**NAC_methylation**: enhancer regions as defined by DNA methylation only (non-activated CD4^+^ T cells). **NAC.Methylation.prediction**: prediction rate of enhancer regions as defined by DNA methylation only divided by enhancer regions as defined by histone marks only (non-activated CD4^+^ T cells), **overlap.NAC.AC**: number of DNA methylation predicted regions that are overlapping between activated and non-activated CD4^+^ T cells. **AC_histones**: enhancer regions as defined by histone marks only (activated CD4^+^ T cells). **AC_methylation**: enhancer regions as defined by DNA methylation only (activated CD4^+^ T cells). **AC.Methylation.prediction**: prediction rate of enhancer regions as defined by DNA methylation only divided by enhancer regions as defined by histone marks only (activated CD4^+^ T cells), **% switch**: the percentage of DNA methylation predicted regions that are found in CD4^+^ T cells switching from a non-activated to an activated state. **OFF(NAC).TO.ON(AC)**: the number of DNA methylation predicted regions that are found in activated CD4^+^ T cells (absent in non-activated). **ON(NAC).TO.OFF(AC)**: the number of DNA methylation predicted regions that are found in non-activated CD4^+^ T cells (absent in activated). **OFF(NAC).TO.ON(AC) %**: the percentage of DNA methylation predicted regions that are found in activated CD4^+^ T cells (absent in non-activated). **ON(NAC).TO.OFF(AC) %**: the percentage of DNA methylation predicted regions that are found in nonactivated CD4^+^ T cells (absent in activated). *The % within brackets corresponds to the % of verified states predicted by methylation based on the histone gold standard*.

| Enhancer | NAC_histones | NAC_methylation | NAC.Methylation.prediction | overlap.NAC.AC |
| --- | --- | --- | --- | --- |
| Active | 6189 | 5848 | 0.94 | 3932 |
| Poised | 7778 | 6822 | 0.88 | 123 |
| promoter_proximal | 245 | 222 | 0.91 | 163 |
| putative | NA | 789 | NA | 680 |
| Total | 14212 | 13681 | 0.91 | 4898 |

| Enhancer | AC_histones | AC_methylation | AC.Methylation.prediction | % switch |
| --- | --- | --- | --- | --- |
| Active | 9912 | 9184 | 0.93 | 0.67 |
| Poised | 274 | 239 | 0.88 | 0.51 |
| promoter_proximal | 565 | 522 | 0.92 | 0.73 |
| Putative | NA | 1052 | NA | 0.86 |
| Total | 10751 | 10997 | 0.91 | 0.69 |

| Enhancer | OFF(NAC).TO.ON(AC)* | ON(NAC).TO.OFF(AC)* | OFF(NAC).TO.ON(AC) % | ON(NAC).TO.OFF(AC) % |
| --- | --- | --- | --- | --- |
| Active | 5252 (93%)* switch to active in activated CD4+ T cells | 1916 (97%)*switch to active in non-activated CD4+ T cells | 86 | 22 |
| Poised | 116 (98%)* switch to poised in activated CD4+ T cells | 6699 (88%)* switch to poised in non-activated CD4+ T cells | 2 | 76 |
| promoter_proximal | 359 (94%) | 59 (94%) | 6 | 1 |
| latent (methylation) | 372 | 109 | 6 | 1 |
| Total | 6099 | 8783 | 100 | 100 |

Given the accuracy (ACC = 0.90), sensitivity (TPR = 0.91) and specificity (SPC = 0.99), calculated over both activated and non activated enhancers, this is statistically significant result (hypergeometric test, P < 0.001), which implies that DNAm oscillations alone may be used as a predictor of enhancer regions, and that these regions are defined by the methylome irrespective of the activation state.

Based on the validation using histone marks, we also found that 67% of active enhancers defined by DNA-methylation were found in both activated and non-activated states. We hypothesize that the latter regions may participate in changing toward an activated phenotype, as well defining CD4^+^ T cell lineage. Analogously, based on histone marks validation, we found that 51% of methylation-defined **poised** enhancers were found in both states, as well as 73% of promoter proximal regions. At the same time 86% of regions defined by methylation alone (not confirmed by histone marks) were found in both activated and non-activated CD4^+^ T cell methylomes. Based on the DNAm oscillation prediction (**Table 1**), 5596 regions switch from inactive to active in activated CD4+ T cells, 93% of which (5252) are predicted by methylation. Likewise, 7622 switch to poised in non-activated CD4^+^ T cells, of which 6699 (88%) are predicted by methylation. Also, 372 regions were marked by DNAm only prior to activation and 109 regions were marked by DNAm only prior to deactivation. According to our methylome analysis of CD4^+^ activation, 86% of regions which are switched on (from poised to active) upon activation are predicted to become active enhancers, and 76% of the regions which are switched off (from active to poised) are predicted to become poised enhancers in activated CD4^+^ T cells. Taken together, these results lead us to hypothesize that methylation alone can be used to predict the location of latent enhancers, previous to their activation.

### Machine learning models support DNAm as predictive of enhancer location

Using enhancers with strong signal as defined by BLUEPRINT segmentations and ENSEMBL regulatory build as reference set, we used an array of machine learning algorithms to test the hypothesis that DNAm oscillations may be informative for predicting enhancers using genome-wide data for each of the predictive categories at the resolution used for the enrichment analyses (**Methods**). As input, we used the data employed for the third analysis (BLUEPRINT segmentations integrated in conjunction with the ENSEMBL Regulatory Build) where the rows of the matrix represent genomic coordinates, and columns represent features. The response variable to be predicted was a class label of 0 (negative controls) and 1 (enhancers). To summarize model performance, we estimated the % of times in which each feature was estimated to be informative for predicting enhancers. In the first instance, we built a partial least squares regression model with recursive feature elimination. In this analysis, we found DNAm oscillations to be selected as an informative feature for predicting enhancers 100% of the times, with histone marks also being very informative (**Supplementary Table 7**). Crucially, H3K4me3 marks are useful for sorting promoters from enhancers when using models agnostic to gene location. In this context, it is important to note that absence of a mark can be just as informative as a presence, as H3K36me3 marks are inversely related to enhancer location (r = −0.1, p < 0.001). Collectively, all features are informative, as error decreases when the number of features increases, with four features being sufficient (**Supplementary Figure 8**). ROC analysis showed the model to be of high performance (AUC = 0.90, **Supplementary Figure 9**).

We estimated the informative nature of DNAm oscillations to be predictive of enhancer locations based on additional machine learning algorithms, including generalized linear models (**Methods**) in conjunction with forward feature selection (AUC = 0.92). This analysis highlighted the predictive value of DNAm oscillations where it was always selected as the more informative feature (**Supplementary Figure 10**), here a total of four features were also found to be best suited for predicting enhancers (**Supplementary Figure 11**).

The results of the elastic nets model (**Methods**) also ranked DNAm oscillations to be predictive, though this was only run on a subset of the data (AUC =0.88, **Supplementary Table 8**). All in all, DNAm oscillations were found to be informative by all the methods employed (**Supplementary Table 7**). Interestingly, H3K4me3, H3K4me1 and target regions from PCHi-C regions are less informative than the other features, when taken individually (**Supplementary Table 7**).

In order to include positive and negative controls, and account for additional features related to enhancer prediction and include other features related to DNA methylation, we repeated the analysis by adding four more features, namely CpG islands (CGI), low-methylated regions (LMR) and unmethylated regions (UMR) using an established DNAm segmentation algorithm based on Hidden Markov Models (**Methods**) and DNaseI Hypersensitivity Clusters in 125 cell types from ENCODE (DNAase_cluster). Given the new input variables, this 10-feature analysis confirmed the overall ranking and informative levels of individual features as measured by the three models (**Supplementary Table 7**). In addition, we estimated the overall informative levels of the DNAase_cluster to be at 73.3%, CGI at 41.3%, UMR at 20% and LMR at 0% (**Supplementary Table 7**).

## Discussion

In eukaryotes, enhancers control the activation of gene expression. Importantly, the activity of enhancers can be restricted to a cell type, a time point, or a specific condition [6]. For example, the activation of CD4^+^ T cells is a pronounced epigenetic and transcriptional event that has been described after four hours [13]. In this context, understanding the dynamic interaction between genomic and epigenomic signals is key for identifying such regions and inferring their function [6,8]. While there is no single enhancer mark, histone modifications and chromatin structure have long been associated with the search of candidate enhancers [6]. These regions can be defined as elements that increase transcription level independently of their orientation, position and distance to a promoter and can be characterized with histone modification profiles, open chromatin information, transcription factor binding sites and other types of data with increased accuracy in a cell-type specific context [7]. Long-range interactions between promoters and enhancers have also been identified using chromosome conformation capture techniques, such as promoter capture Hi-C (PCHi-C). For example this method has recently been applied to 17 human primary haematopoietic cell types, including the ones described here [29]. In our work, we use target (enhancer) regions from this study in conjunction with a whole compendium of BLUEPRINT histone mark data matched to whole genome bisulfite sequencing methylomes in multiple blood cell types to estimate the power of DNAm oscillations to predict enhancers.

Using the above mentioned data types as reference sets, we show that DNA methylation is a genomic signal [8] and epigenomic marker whose spatial patterns (DNAm oscillations, as frequency of harmonics) [14] can be used to infer the location of enhancer regions. Having observed that increased frequency of harmonics can be used to discriminate different types of genomic regions in a test set (**Figure 1**), we observed a 30-fold difference in oscillatory patterns between background and enhancers based on multiple analyses (**Supplementary Table 2**, **Figure 2**).

Having established the enrichment of DNAm in enhancers based on reference regions identified with other data, we set out to study the percentage of recovery of enhancers using DNAm oscillations alone. We were able to recover 90% of enhancers of strong signal using DNAm oscillations alone across all 9 cell types (**Figure 3**). Using multiple machine learning models (**Methods, Results**), we estimated the predictive power of DNAm oscillations for estimating enhancer types when compared to the other datasets employed in this study. DNAm oscillations were found to be informative for estimating enhancer locations alongside histone marks (**Supplementary Table 7**), with DNAm oscillations were found to be the most consistently informative individual feature across all models.

The power of DNAm oscillation in predicting enhancers was confirmed when we tested the hypothesis that DNA methylation alone is sufficient for inferring the genome regulatory backbone. In this case, CD4^+^ T cell activation methylomes were generated in the context of a parallel study [13], where the experimental design was such that within four hours, the activation of CD4^+^ T cells gave rise changes in the activity of regulatory elements and in the transcription of enhancer RNAs that corresponded to changes in the expression of their interacting target genes identified by PCHi-C [12,13]. In this setting, we used enhancer regions defined in [12,13] as a reference set, and our DNAm data supports the evidence that 90% of activated enhancers can be recovered using DNAm oscillations alone (**Table 1**).

Enhancer properties such as cell specificity, redundancy and plasticity [1] are the background against which we set our hypothesis. Indeed, we hypothesize DNAm oscillations to be a cell-specific long-term [8] marker laying out the epigenetic landscape against which multiple chromatin events may occur, with DNA binding proteins reading the CpG signal [8], and in turn activating or repressing enhancer regions. Our results on the methylomes of activated and non-activated CD4^+^ T cells indicate that DNAm oscillations exist in both states, irrespective of activation, whereas histone marks may become detached in non-activated states. Latent were originally identified upon activation by stimulation of differentiated cells [3]. As such, DNAm oscillations could be used to glean the location of latent enhancers even before the activation of such regions.

This work highlights the reproducibility of DNAm oscillations as predictor of enhancers across multiple cell types, even in the context of cell specificity and variability among samples. Our results indicate that these regions are identifiable by DNA methylation alone irrespective of the epigenetic state. Based on this, it is possible to hypothesize that DNA methylation may act as an epigenetic backbone which marks regions which are later used as genome occupancy regions for histone marks at any stage of development. This work is the first to suggest that DNA methylation alone may be used to define enhancers, based on its spatial oscillatory patterns, and not dynamic CpG regions or differentially methylated regions. Also, we suggest that this epigenomic mark defining regulatory regions is stable across cell lineage and activation.

## Acknowledgments

We gratefully acknowledge the participation of all NIHR Cambridge BioResource volunteers. We thank the Cambridge BioResource staff for their help with volunteer recruitment. We thank members of the Cambridge BioResource SAB and Management Committee for their support of our study and the National Institute for Health Research Cambridge Biomedical Research Centre for funding. KD is funded as a HSST trainee by NHS Health Education England. MF is supported by the BHF Cambridge Centre of Excellence [RE/13/6/30180]. Research in the Ouwehand laboratory is supported by EU-FP7 project BLUEPRINT (282510) and by program grants from the National Institute for Health Research (NIHR, http://www.nihr.ac.uk); and the British Heart Foundation under numbers RP-PG-0310-1002 and RG/09/12/28096 (http://www.bhf.org.uk). The laboratory receives funding from the NHS Blood and Transplant for facilities.

This work was funded by the JDRF (9-2011-253), the Wellcome Trust (089989, 091157, 107881), the UK Medical Research Council (MC_UP_1302/5), the UK Biotechnology and Biological Sciences Research Council (BB/J004480/1) and the National Institute for Health Research (NIHR) Cambridge Biomedical Research Centre. The research leading to these results has received funding from the European Union’s 7^th^ Framework Programme (FP7/2007-2013) under grant agreement no.241447 (NAIMIT). The Cambridge Institute for Medical Research (CIMR) is in receipt of a Wellcome Trust Strategic Award (100140).

RH acknowledges Dangoor Education for their support towards this work. S.C.H., M.G., I.G.G. and H.G.S. were supported by EU-FP7 project BLUEPRINT (282510). E.L. and S.B. were supported by EU-FP7 projects EpiTrain (316758), EpiGeneSys (257082) and BLUEPRINT(282510), the Wellcome Trust (99148) and a Royal Society Wolfson Research Merit Award (WM100023).

